# Detecting intragenic *trans*-splicing events with hybrid transcriptome sequencing in cancer cells

**DOI:** 10.1101/2022.04.21.489006

**Authors:** Yu-Chen Chen, Chia-Ying Chen, Tai-Wei Chiang, Ming-Hsien Chan, Michael Hsiao, Huei-Mien Ke, Isheng Jason Tsai, Trees-Juen Chuang

**Affiliations:** Genomics Research Center, Academia Sinica, Taipei, Taiwan.; Biodiversity Research Center, Academia Sinica, Taipei, Taiwan.

## Abstract

*Trans*-splicing can generate non-co-linear (NCL) transcripts that consist of exons in an order topologically inconsistent with the corresponding DNA template. Detecting *trans*-spliced RNAs (ts-RNAs) may be interfered by false positives from experimental artifacts, circular RNAs (circRNAs), and genetic rearrangements. Particularly, intragenic ts-RNAs, which are derived from separate precursor mRNA molecules of the same genes, are often mistaken for circRNAs through analyses of high-throughput transcriptome sequencing (RNA-seq) data. In addition, the biogenesis and function of ts-RNAs remain elusive. Here we developed a bioinformatics pipeline, NCLscan-hybrid, with the integration of long and short RNA-seq reads to minimize false positives and identify intragenic ts-RNAs. We utilized two features of long reads, out-of-circle and rolling circle, to distinguish intragenic ts-RNAs from circRNAs. We also designed multiple experimental validation steps to examine each type of false positives and successfully confirmed an intragenic ts-RNA (ts-ARFGEF1) in breast cancer cells. On the basis of ectopic expression and CRISPR-based endogenous genome modification experiments, we confirmed that ts-ARFGEF1 formation was significantly dependent on the reverse complementary sequences in the flanking introns of the NCL junction. Subsequent *in vitro* and *in vivo* experiments demonstrated that ts-ARFGEF1 silencing can significantly inhibit tumor cell growth. We further showed the regulatory role of ts-ARFGEF1 in p53-mediated apoptosis through affecting the PERK/eIF2a/ATF4/CHOP signaling pathway in breast cancer cells. This study thus described both bioinformatics procedures and experimental validation steps for rigorous characterization of transcriptionally non-co-linear RNAs, expanding the discovery of this important but understudied class of RNAs.

## Introduction

Precursor mRNA (pre-mRNA) splicing is an important step in generating mature mRNA, which can occur either in *cis* or *trans*. *Cis*-splicing occurs within a single pre-mRNA, whereas *trans*- splicing occurs between two or more separate pre-mRNAs arising from the same genes (intragenic *trans*-splicing) or different genes (intergenic *trans*-splicing) (Horiuchi and Aigaki 2006; Gingeras 2009). Thus, *trans*-splicing often forms non-co-linear (NCL) transcripts that consist of exons in an order topologically inconsistent with the corresponding DNA template. Regarding the transcripts generated during the transcriptional process, however, an observed intragenic NCL event may also arise from *cis*-backsplicing (Yu et al. 2014; Chen et al. 2015), which form non-polyadenylated and RNase R-resistant circular molecules with a covalently closed continuous loop structure (Chen 2016; Li et al. 2018) (the so-called circular RNA or “circRNA”). Analyses of high-throughput transcriptome sequencing (RNA-seq) data from total RNAs without poly(A)-selection have uncovered a tremendous number of intragenic NCL transcripts in diverse species (Glazar et al. 2014; Dong et al. 2018; Ji et al. 2019; Chen 2020; Wu et al. 2020). In spite of the fact that the majority of the detected intragenic NCL junctions generally originate from circRNA products, genome-wide comparisons of poly(A)-selected and poly(A)-depleted RNA-seq data suggested that a considerable percentage of the detected intragenic NCL junctions were derived from ts-RNA products (Chuang et al. 2016; Chuang et al. 2018). In fact, a few intragenic NCL RNAs observed in human cell lines were experimentally confirmed to be generated by *trans*-splicing rather than *cis*-backsplicing (Takahara et al. 2000; Flouriot et al. 2002; Wu et al. 2014; Yu et al. 2014). Although the function of most intragenic ts-RNAs remains elusive, some human intragenic ts-RNA events were confirmed to be expressed across primates and rodents (Takahara et al. 2002; Yu et al. 2014), indicating their evolutionary significance. Our previous study showed that the expression patterns of ts-RNAs and their cognate co-linear isoforms were not correlated exactly with each other (Wu et al. 2014), suggesting that they could participate in different biological functions. In fact, an intragenic ts-RNA (i.e., tsRMST), not its corresponding co-linear counterpart, was demonstrated to serve as an important role in regulating the pluripotency maintenance of human embryonic stem cells (Wu et al. 2014). In contrast with widely characterized circRNAs, intragenic ts-RNAs remain largely unexplored so far.

There are several challenges in investigation of intragenic ts-RNAs. First, like other types of NCL events (e.g., circRNAs), a ts-RNA event is generally detected according to short reads of Second Generation Sequencing that span an NCL junction. Such a short-read strategy is often hampered by misalignments (due to genome variants or sequencing errors) or ambiguous alignments (Treangen and Salzberg 2011; Gao et al. 2015; Chen and Chuang 2019a) (due to numerous repeated sequences or paralogous genes in the human genome). Second, as mentioned above, it is difficult to effectively distinguish intragenic ts-RNAs from circRNAs on the basis of short RNA-seq reads. It is known that circRNAs are generally non-polyadenylated but ts-RNAs are not (Salzman et al. 2012; Memczak et al. 2013; Guo et al. 2014; Suzuki and Tsukahara 2014; Yu et al. 2014; Zhang et al. 2014; Chen et al. 2015; Chen and Yang 2015). However, circRNAs may be detected in poly(A)-selected RNA-seq data because of incomplete depletion of poly(A)-tailed RNAs (Wang et al. 2014; Gao et al. 2015; Chuang et al. 2016; Nair et al. 2016). In some cases, intragenic ts-RNAs and circRNAs may share the same NCL junctions (Yu et al. 2014; Chuang et al. 2018), further complicating the identification of ts- RNAs. Third, experimental artifacts such as template switching events frequently emerge in cDNA products (Cocquet et al. 2006; Houseley and Tollervey 2010) and are often misinterpreted as transcriptional NCL events (McManus et al. 2010). Such artifacts cannot be easily detected by merely *ab initio* controlling for the number of supported RNA-seq reads spanning the NCL junctions (McManus et al. 2010; Wu et al. 2014; Yu et al. 2014). Fourth, genetic rearrangement events occurring at the DNA level can also form NCL events and masquerade as ts-RNAs (Gingeras 2009), arising another challenge to detecting true ts-RNAs. Fifth, it remains unclear whether ts-RNA products are merely side-products of imperfect pre-mRNA splicing or can play a role in mediating the expression of their corresponding co-linear host genes, although the host genes are the source of ts-RNA isoforms. Finally, the biogenesis of intragenic ts-RNA isoforms remains unclear. Although ts-RNA formation was reported to be associated with reverse complementary sequences (RCSs) residing in the introns flanking the NCL junctions (Dixon et al. 2007; Chuang et al. 2018), endogenously experimental evidence supporting the model is still lacking.

Single-molecule long reads generated from Third Generation Sequencing technologies such as Pacific Biosciences (PacBio) and Oxford Nanopore Technologies (ONT) offer a unique advantage to recognize NCL transcript isoforms. The long-read sequencing platforms can generate tens of thousands of bases in length. With the support of such an ultra-long length, we can minimize the possibility of short-read-identified NCL junctions that were derived from alignment ambiguity. In addition, a non-co-linearly matched long read may span a short-read- identified NCL junction and map outside the predicted circle of a possible circRNA isoform, making long-read RNA-seq a good indicator for distinguishing ts-RNAs from circRNAs. In this study, we presented a pipeline, NCLscan-hybrid, to identify intragenic ts-RNAs through the hybrid sequencing strategy integrating Illumina short reads and PacBio long reads from MCF-7 human breast cancer cell line. To confirm the identified intragenic ts-RNAs, we also designed multiple experimental validation steps to detect false positives from experimental artifacts, circRNAs, or genetic rearrangements. We further designed an ectopic expression system and endogenous genome modification experiments to confirm the requirement of RCSs in the flanking introns of NCL junctions for the formation of ts-RNAs. Finally, we experimentally examined the biological function of the selected ts-RNA and its regulatory role in breast cancer development. Our findings provided rigorous *in silico* procedures and experimental validation steps to characterize intragenic NCL events and showed further insight into the potential roles of intragenic ts-RNAs in breast cancer development.

## Results

### A bioinformatic pipeline for detecting intragenic ts-RNAs

To detect intragenic ts-RNAs, we developed the pipeline (“NCLscan-hybrid”) capable of integrating both short- and long RNA-seq reads (Fig. 1A). We extracted Illumina short and PacBio long polyadenylated RNA-seq data of MCF-7 from the ENCODE project (Bernstein et al. 2012; Djebali et al. 2012) and the PacBio dataset, respectively (Table 1). NCLscan-hybrid first employed NCLscan (Chuang et al. 2016) to search for all possible intragenic ts-RNA candidates according to the short reads. Of note, NCLscan is a short read-based detector, which was demonstrated to be an accurate NCL event detector adept at effectively eliminating alignment artifacts (Chuang et al. 2016). A total of 1,066 intragenic NCL junctions were detected (Supplemental Table S1). NCLscan-hybrid generated a pseudo sequence by concatenating the exonic sequences flanking the NCL junction for each detected NCL junction and then aligning all PacBio long reads against the pseudo sequence (Fig. 1B). Only the NCL events with the support of PacBio long reads were retained. Next, each PacBio long read that supported an NCL junction was split into two segments at the NCL junction, of which each segment was aligned against the reference genome. To minimize false positives from ambiguous alignments with an alternative co-linear explanation or multiple hits, we only considered the NCL events that satisfied the two rules simultaneously (Fig. 1B and Methods). First, both segments of a supporting PacBio long read were uniquely matched to the same gene loci. Second, integration of both mapped segments can span the specified NCL junction. After that, 42 NCL events supported by both Illumina short reads and PacBio long reads were retained. Since a few circRNA events may persist during poly(A) selection (Gao et al. 2015; Chuang et al. 2016), the detected NCL events may originate from residual circRNAs. We utilized the supporting PacBio long reads to distinguish ts-RNA events from circRNA ones. We considered the NCL events supported by at least one long read that spanned the NCL junctions (e.g., the Exon4-Exon3 junction; Fig. 1B) and mapped outside the potentially predicted circle sequences of the circRNA isoforms (e.g., Exon2-Exon3-Exon4-Exon3-Exon4 or Exon3-Exon4-Exon3- Exon4-Exon5) as ts-RNA events. Such a long read was designated as an “out-of-circle read”. A total of 18 ts-RNA candidates with the support of out-of-circle reads were retained. An example of a ts-RNA candidate (i.e., ts-ARFGEF1) supported by 7 out-of-circle reads was shown in Figure 1C. In addition to the 18 ts-RNAs, we found three NCL events with supporting long reads that captured the whole predicted circle sequence and crossed the NCL junction twice (see Fig. 1B). In such a case, the long read was split into three segments at the NCL junction when aligning the long read against the pseudo sequence, but did not map outside the potentially predicted circle sequences of the circRNA isoforms. Such a long read was designated as a “rolling-circle read”. The NCL events supported by rolling-circle reads were regarded as circRNAs. An example of rolling-circle read-supported NCL events (i.e., circCDYL) was given in Figure 1D. circCDYL has been experimentally confirmed (Wei et al. 2020) and demonstrated to be associated with breast cancer progression (Liang et al. 2020), again supporting the abovementioned notion that there were residual circRNAs in poly(A)- selected RNA-seq data due to incomplete depletion of poly(A)-tailed RNAs (Wang et al. 2014; Gao et al. 2015; Chuang et al. 2016). This observation also showed that NCLscan-hybrid was able to distinguish ts-RNAs from circRNAs with presence of long reads. Finally, to check whether the identified ts-RNAs were derived from genetic rearrangements, we downloaded whole genome sequencing data (WGS) of MCF-7 (Table 1) and performed INTEGRATE (Zhang et al. 2016) to detect structural variants. Indeed, we did not detect structure variants (e.g., duplications or breakpoints; Supplemental Table S1) in the flanking introns of the NCL junctions for the 18 ts-RNA candidates.

**Figure 1.**
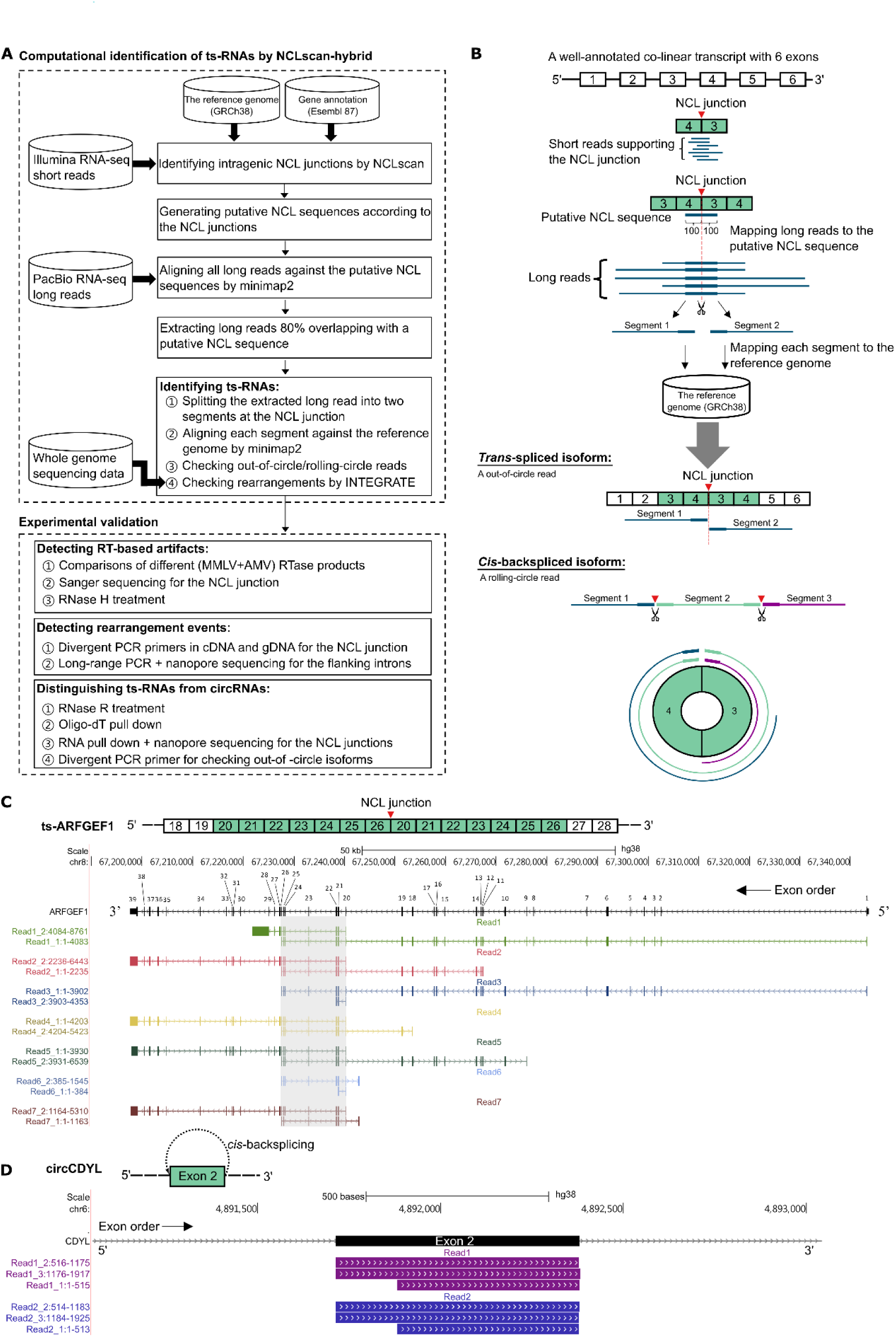
Identification of intragenic ts-RNAs. **A** Flowchart of NCLscan-hybrid identification and subsequent experimental validations for ts-RNAs. **B** Schematic illustrations of NCLscan-hybrid processes. NCLscan-hybrid utilized out-of-circle and rolling-circle reads to distinguish between intragenic ts-RNAs (*trans*-spliced isoform) and circRNAs (*cis*-backspliced isoform). **C, D** Examples of the identified intragenic ts- RNA candidate with the supporting out-of-circle reads (ts-ARFGEF1, **C**) and circRNA candidates with the supporting rolling-circle read (circCDYL, **D**).

**Table 1.**
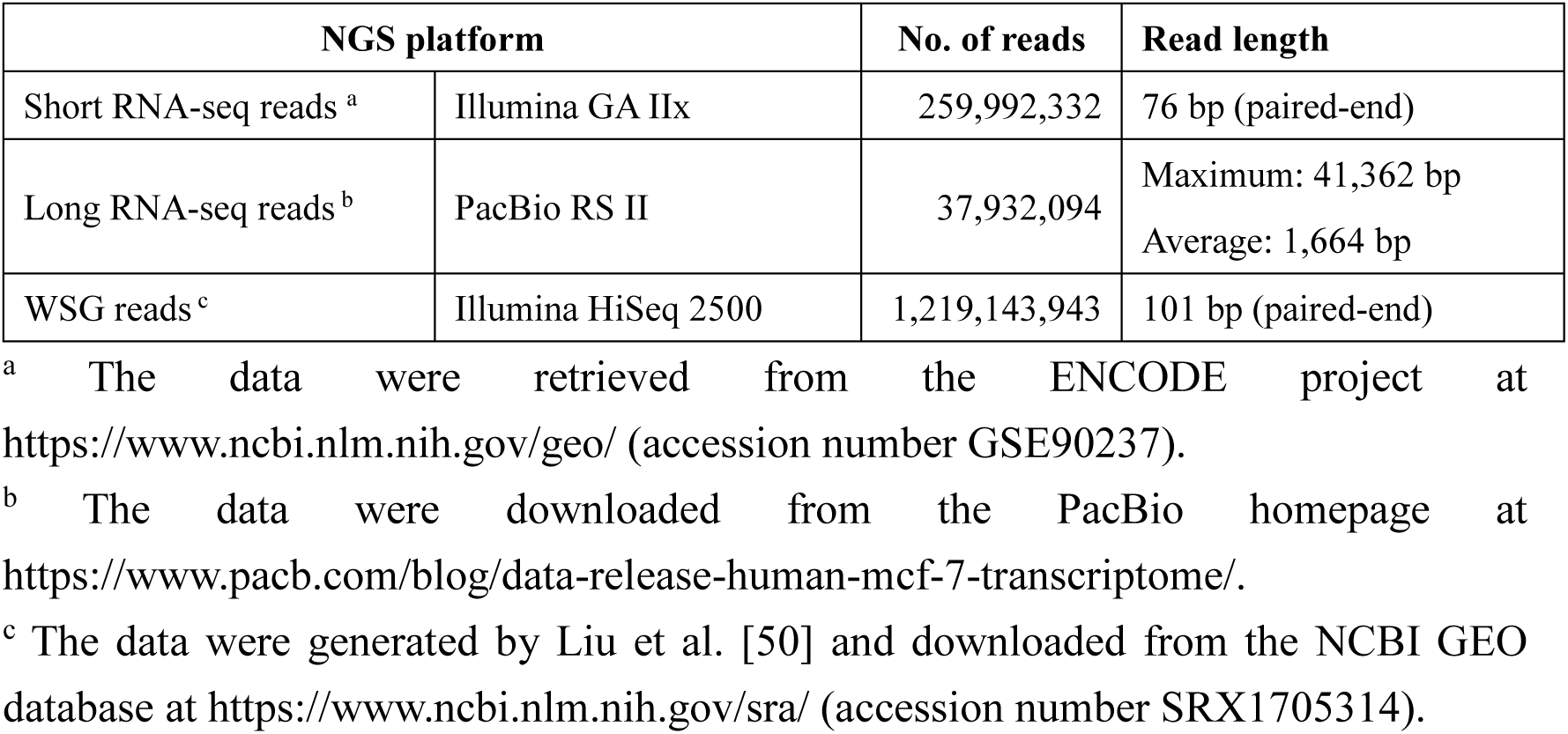
MCF-7 NGS data used in this study.

| NGS platform |  | No. of reads | Read length |
| --- | --- | --- | --- |
| Short RNA-seq reads <sup>a</sup> | Illumina GA IIx | 259,992,332 | 76 bp (paired-end) |
| Long RNA-seq reads <sup>b</sup> | PacBio RS II | 37,932,094 | Maximum: 41,362 bp<br>Average: 1,664 bp |
| WSG reads <sup>c</sup> | Illumina HiSeq 2500 | 1,219,143,943 | 101 bp (paired-end) |

<sup>a</sup> The data were retrieved from the ENCODE project at https://www.ncbi.nlm.nih.gov/geo/ (accession number GSE90237).
<sup>b</sup> The data were downloaded from the PacBio homepage at https://www.pacb.com/blog/data-release-human-mcf-7-transcriptome/.
<sup>c</sup> The data were generated by Liu et al. [50] and downloaded from the NCBI GEO database at https://www.ncbi.nlm.nih.gov/sra/ (accession number SRX1705314).

### Multiple experimental validation steps for detecting potential false positives

We quantified the abundance of the identified ts-RNA candidates using the short and long polyadenylated RNA-seq reads that supported the NCL junctions. For the short RNA-seq reads, we calculated the NCL ratio (Chuang et al. 2018; Chen and Chuang 2019b) of a ts-RNA candidate at the NCL junction on the basis of the number of reads spanning the NCL junction and the number of reads spanning both the co-linearly spliced junctions (the donor and acceptor sites; Fig. 2A). We noted that one of the identified ts-RNA candidates (i.e., ts-ARFGEF1) was featured the highest NCL ratio. We found that the PacBio long reads spanned the NCL junction (Exon26-Exon10) and mapped outside the possible circle sequences (Exon20-Exon21-Exon22- Exon23-Exon24-Exon25-Exon26; see Fig. 1C), supporting the existence of the *trans*-spliced isoform. We then designed a systematic pipeline with multiple experimental validation steps to examine whether this detected ts-RNA event was a false positive derived from (1) RT-based artifact, (2) genetic rearrangement, or (3) circRNA (see also Fig. 1A). First, we examined whether ts-ARFGEF1 was derived from RT-based artifacts such as template switching events (Cocquet et al. 2006; Houseley and Tollervey 2010). It has been reported that comparisons of different RTase products can effectively detect RT-based artificial NCL junctions because RTase-dependent products tend to arise from RT-based artifacts (Houseley and Tollervey 2010; Wu et al. 2014; Yu et al. 2014; Chen et al. 2015; Chuang et al. 2018). We thus performed reverse transcription polymerase chain reaction (RT-PCR) based on Avian Myeloblastosis Virus (AMV)- and Moloney Murine Leukemia Virus (MMLV)-derived RTase in parallel experiments in MCF-7 cells, followed by Sanger sequencing of the RT-PCR amplicons to confirm the NCL junction (Fig. 2B). Our result revealed that the NCL junction of ts-ARFGEF1 was RTase- independent. After that, we performed RNA-Fluorescence *in situ* hybridization (RNA-FISH) analysis and showed that this NCL event was expressed in the nucleus (Fig. 2C), supporting that this NCL event was not an RT-based artifact. Consistently, the subcellular fractionation analysis also revealed that this NCL event was predominantly expressed in the nucleus (Supplemental Fig. S1). In addition, we designed specific locked nucleic acid (LNA)-modified antisense oligonucleotides (ASO) to target the NCL junction of ts-ARFGEF1 and then treated total RNA from MCF-7 cells with RNase H to digest the transcripts containing the target region at the RNA level (Fig. 2D, top). We performed qRT-PCR analysis to examine the expression of the transcripts containing the NCL junction. Such expression would not be significantly affected by RNase H treatment, if the NCL junction originated from an experimental artifact during RT. We showed that the expression of the transcripts containing the NCL junction significantly decreased after RNase H degradation (Fig. 2D, bottom). The above results thus suggested that the NCL junction of ts-ARFGEF1 was not derived from an RT-based artifact.

**Figure 2.**
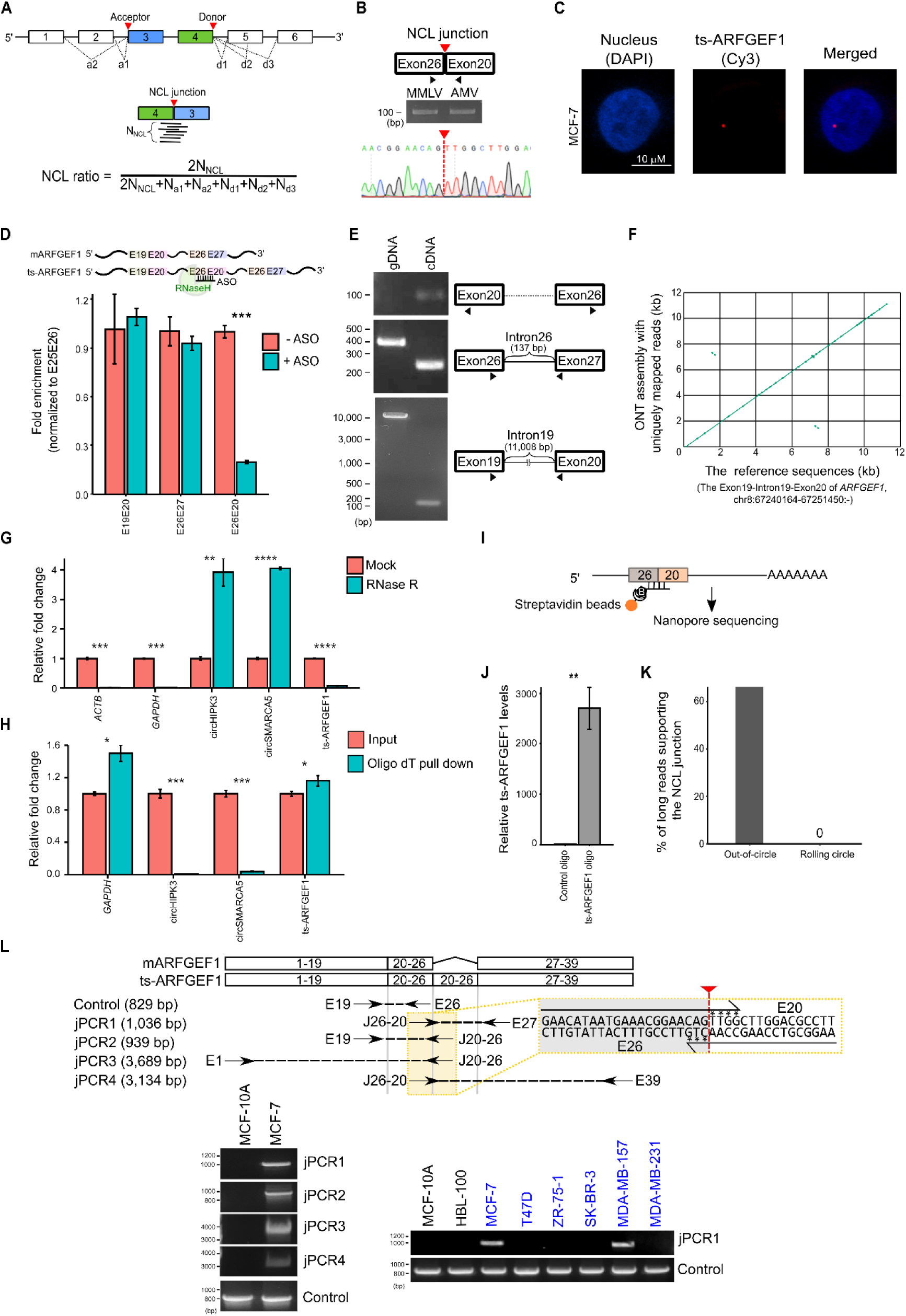
Experimental validations of the selected ts-RNA. **A** Schematic illustration of the evaluation of NCL ratio (top) and two-dimensional scaling screen for the most highly abundant ts-RNA (ts-ARFGEF1) among the identified ts-RNA candidates (bottom). N_NCL_ represents the number of short reads spanning the NCL junction. d_i_ and a_i_ represent the numbers of short reads spanning the corresponding co-linear junctions at the NCL donor and acceptor sites, respectively. **B-L** Experimental examinations of potentially false positives from RT-based artifacts (**B-D**), genetic rearrangements (**E, F**), and circRNAs (**G-L**). **B** Comparison of two different RTase (MMLV and AMV-derived) products of the ts-ARFGEF1 junction (Exon26-Exon20), followed by Sanger sequencing the RT-PCR amplicons for this NCL event in MCF-7. **C** Subcellular location analysis of the ts-ARFGEF1 event. The Cy3 labeled probes were targeted against the ts-ARFGEF1 junction to determine the location of ts-ARFGEF1 by RNA-FISH. **D** qRT-PCR analyses of potential RT-based artifacts by RNase H cleavage of the LNA-modified ASO targeted transcripts. The absence of oligonucleotides served as the control. **E** Detection of structural variants (i.e., duplication/insertion) in Introns 26 and 20 by PCR and long PCR experiments, respectively. **F** Genome alignment between the consensus sequence formed by the generated nanopore reads and the reference genomic sequence of *ARFGEF1* Intron 19. **G, H** qRT-PCR analyses of expression fold changes for the NCL junction of ts-ARFGEF1 in MCF-7 before/after RNase R treatment (**G**) or oligodT pull down (**H**). **I-L** Validation of the existence of the *trans*-spliced isoform. A schematic diagram represented the transcripts pulled down by biotin-labeled oligonucleotides targeting the NCL junction of ts-ARFGEF1, followed by nanopore long-read sequencing (**I**). The pull-down efficiency was shown (**J**). Comparison of out-of-circle and rolling-circle long reads based on the generated nanopore reads was performed (**K**). Four pairs of convergent PCR primers (**L**, top) were designed to confirm the existence of ts-ARFGEF1 using RT-PCR (**L**, bottom left). RT-PCR experiments showed that ts-ARFGEF1 was expressed in multiple breast cancer cell lines (MCF-7 and MDA-MB-157; **L**, bottom right). All error bars represented the means ± standard deviation of three independent experiments. *P* values were determined using two-tailed *t*-test. \**P*<0.05, \*\**P*<0.01, \*\*\**P*<0.001.

Second, we examined whether the NCL event was generated from a genetic rearrangement event. Regarding the two flanking introns of the NCL junction, i.e., the intron between Exon 26 and Exon 27 (Intron 26) and the intron between Exon 19 and Exon 20 (Intron 19) (Fig. 2E), we conducted the following experiments. For Intron 26 (a short intron with 137 bp in length), we performed a PCR experiment, followed by Sanger sequencing, to confirm the intron (length and intronic sequence) in MCF-7 cells (Fig. 2E). Since Intron 19 is a long intron with >11k bp in length, a long PCR experiment was performed to confirm the length of this intron (Fig. 2E). We further used ONT MinION sequencer (Ip et al. 2015) to generate long genomic reads from MCF-7 cells for the sequence of Intron 19. These nanopore reads were assembled and formed a consensus sequence. We then aligned the consensus sequence against the reference sequence of Intron 19 and did not observe duplications or large insertions in this intron (Fig. 2F). We also aligned the nanopore reads against the reference sequence of Exon 26 and confirmed no insertion of Exon 26 in Intron 19. The above experiments thus eliminated the possibility that the NCL junction of ts-ARFGEF1 was derived from a genetic rearrangement event.

Finally, we would like to confirm that the NCL junction of ts-ARFGEF1 was derived from a *trans*-splicing, not a *cis*-backsplicing event. We treated total RNA from MCF-7 with RNase R, which could digest linear RNAs with free terminal ends. qRT-PCR analysis revealed that ts- ARFGEF1 was digested by RNase R treatment, suggesting that this NCL event was not a circRNA (Fig. 2G). Meanwhile, we performed qRT-PCR analysis using purified mRNA with poly(A)-tails as a template and showed that ts-ARFGEF1 was a poly(A)-tailed RNA (Fig. 2H). Next, we designed biotin-labeled oligonucleotides to target the NCL junction of ts-ARFGEF1 and pulled down the transcripts containing the target region directly, followed by TGS sequencing of the pulled-down transcripts using the ONT MinION sequencer (Fig. 2I). The pull-down efficiency was verified by qRT-PCR (Fig. 2J). We then performed NCLscan-hybrid to examine the generated nanopore reads and observed that 66% of the NCL junction-supporting reads were out-of-circle reads, whereas none of those was rolling-circle read (Fig. 2K). Furthermore, we designed four pairs of convergent PCR primers (e.g., jPCR1∼jPCR4; Fig. 2L, top), each of which one primer spanned the NCL junction and the other one was located outside the potentially predicted circle sequence. The RT-PCR experiments showed that the NCL transcript indeed spanned outside the potentially predicted circle sequence, again supporting the existence of the *trans*-spliced isoform (Fig. 2L, bottom left). We also found that the NCL event was expressed in multiple breast cancer cell lines, suggesting that this event was not detected by chance (Fig. 2L, bottom right). The above results thus supported that the NCL junction of ts-ARFGEF1 arose from *trans*-splicing, not *cis*-backsplicing.

### Formation of ts-ARFGEF1

We proceeded to study the mechanism by which ts-ARFGEF1 was formed. One possible scenario is that RCSs can mediate the formation of ts-ARFGEF1. We first detected an RCS pair (Fig. 3A) in the flanking introns of the NCL junction of ts-ARFGEF1 by aligning between sequences of Intron 19 and Intron 26. We then examined whether the RCS can trigger ts- ARFGEF1 biogenesis using the ectopic expression system and the endogenous genome modification experiment, respectively. For the ectopic expression experiment, we separately inserted the regions containing Exon26-Intron26-Exon27 (wild-type) and Exon26- Intron26mut-Exon27 (with the mutant RCS in Intron 26) into the pFLAG-CMV2 vectors (Fig. 3A, top). These two types of expression vectors were individually transfected into MCF-7 cells. We then used the primer located at FLAG along with the primer located at the ts-ARFGEF1 junction (i.e., Exon26-Exon20) to monitor the ts-ARFGEF1 expression. Indeed, the FLAG- Exon26-Exon20 chimera products were observed in cells transfected with wild-type vector but not observed in those transfected with the vector of the mutant RCS (Fig. 3B). This result revealed that the RCS in Intron 26 was required for the formation of ts-ARFGEF1.

**Figure 3.**
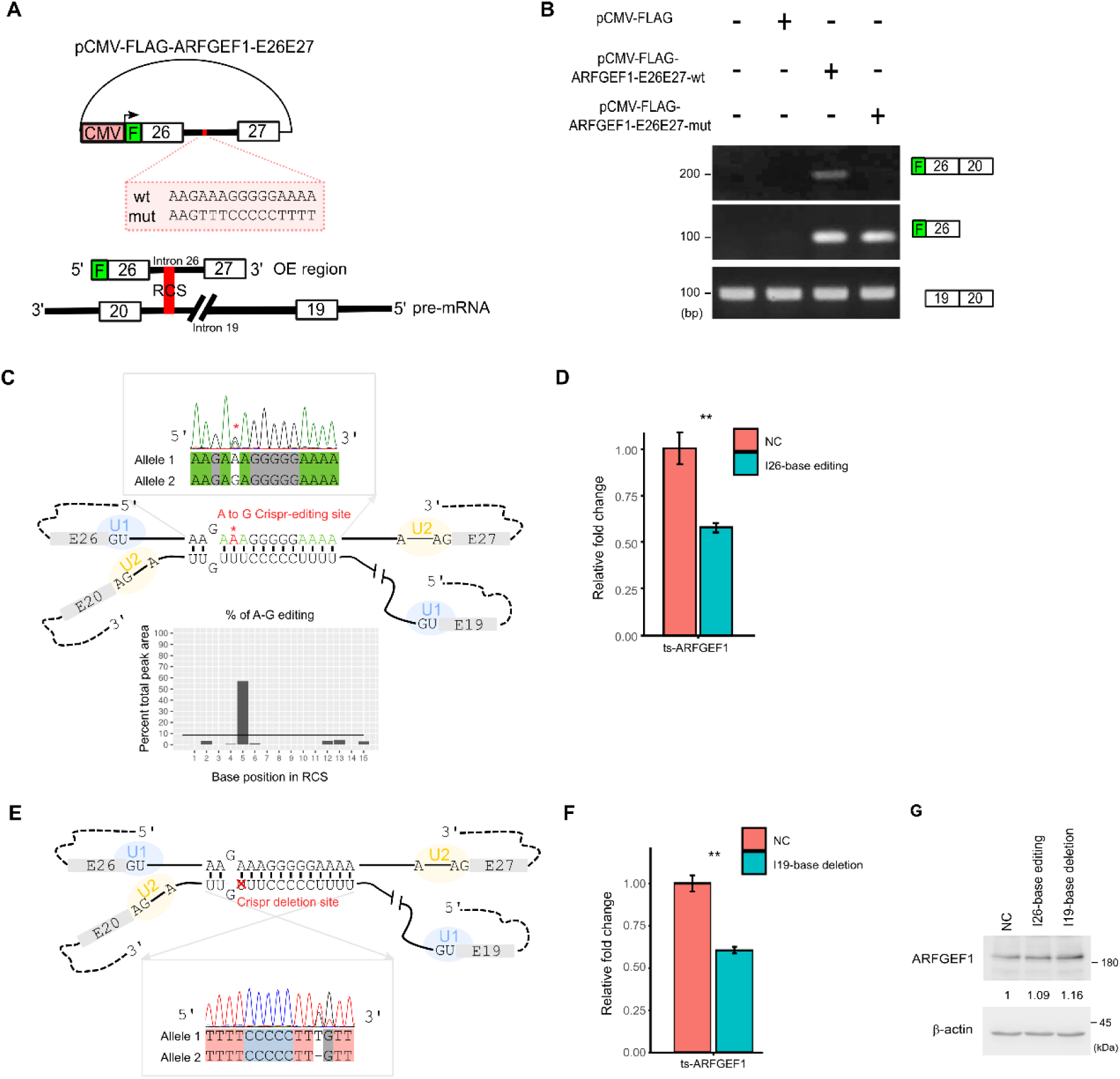
Validation of ts-ARFGEF1 formation with an RCS pair across the flanking introns of the NCL junction. **A** Schematic diagram of the pCMV vector with wild-type and mutant genomic sequences for recapitulation of the FLAG-Exon26-Exon20 chimera products. The genomic regions containing Exon 26, Intron 26, Exon 27, and a FLAG sequence tagged at the 5’ end of Exon 26 (wild-type) and the mutant construct with the mutant RCS were cloned into the pCMV-FLAG vectors, respectively. OE, overexpression. wt, wild-type. mut, mutant. **B** RT-PCR analysis for the expression of FLAG-exon26-exon20 chimera products in MCF-7 cells transfected with different expression plasmids (wild-type, mutant, and empty vectors). The co-linear transcript product with *ARFGEF1* Exons 19 and 20 was used as an internal control. **C, D** Sanger sequencing (**C**) and qRT-PCR analysis (**D**) of CRISPR/Cas9-based genome editing for the RCS in Intron 26 in MCF-7 cells. One nucleotide of the RCS (the position 5 of Allele 2; **C**, top) in Intron 26 was edited from A to G, in which the efficiency of the A-to-G editing was evaluated and plotted by EditR[53] (**C**, bottom). **E, F** Sanger sequencing (**E**) and qRT-PCR analysis (**F**) of high-fidelity CRISPR/Cas9-based genome deletion for the RCS in Intron19 in MCF-7 cells. One nucleotide of the RCS (the position 12 of Allele 2; **E**, bottom) in Intron 19 was deleted. **G** Western blot analysis representing the effects of the CRISPR-based editing and deletion on the protein level of the cognate co-linear form (ARFGEF1). NC, negative control. Error bars represented the means ± standard deviation of three independent experiments. *P* values were determined using two-tailed *t*-test. \*\**P*<0.01.

For the endogenous genome modification experiment, we utilized CRISPR/Cas9 to disrupt the RCS pair within MCF-7 cells. First, we performed CRISPR/Cas9-based genome editing technology to target the RCS in Intron 26 and successfully edited one allele of the RCS (the position 5 of allele 2 was edited from A to G; Fig. 3C). Our result revealed that the expression of ts-ARFGEF1 was significantly reduced when the RCS was edited (Fig. 3D). Second, we employed a high-fidelity CRISPR/Cas9 system to target the RCS in Intron 19 and successfully delete one nucleotide of one allele of the RCS (the position 12 of allele 2; Fig. 3E). Indeed, the deletion can remarkably decrease the expression of ts-ARFGEF1 (Fig. 3F). Of note, Western blot analysis showed that the above CRISPR-based editing/deletion did not affect the protein level of the cognate co-linear form (ARFGEF1) (Figs. 3G). Taken together, these results demonstrated the requirement of the RCS pair in the flanking introns of the ts-ARFGEF1 junction for ts-ARFGEF1 biogenesis.

### Function of ts-ARFGEF1

We observed that ts-ARFGEF1 seemed to be specifically expressed in two types of breast cancer cell lines (MCF-7 and MDA-MB-157), but not expressed in non-tumorigenic epithelial breast cell lines (MCF-10A and HBL-100) (see Fig. 2L). This prompted us to investigate the biological function of ts-ARFGEF1 in breast cancer cells. We designed locked nucleic acid-modified antisense oligonucleotides (LNA ASOs) to target the ts-ARFGEF1 junction and silence ts-ARFGEF1 in MCF-7 and MDA-MB-157 cells (Fig. 4A). Western blot analysis showed that the knockdown did not affect the protein level of ARFGEF1 (Fig. 4B). We then performed the following three experiments to examine the effects of ts-ARFGEF1 knockdown on MCF-7 and MDA-MB-157 cells. First, the Cell Counting kit-8 (CCK-8) assay indicated that the knockdown significantly suppressed cell proliferative ability in MCF-7 and MDA-MB-157 (Fig. 4C). Second, the colony formation assay showed that the number and size of cell colony formation were markedly reduced after ts-ARFGEF1 knockdown (Fig. 4D and 4E). Third, the flow cytometry results demonstrated that ts-ARFGEF1 knockdown significantly promoted the apoptosis rate of both MCF-7 and MDA-MB-157 cells (Fig. 4F and 4G). We assessed the protein-coding capacity of ts-ARFGEF1 and asked whether the phenotypes observed above were derived from the coding products of ts-ARFGEF1. The predicted ts-ARFGEF1 protein sequence and the cognate ARFGEF1 protein sequence shared partial protein sequence (from the coding start site to Exon 26 of ARFGEF1; Supplemental Fig. S2). We presumed that these two protein-coding sequences used the same reading frame and then performed the Western blot analysis based on the anti-N-terminal region of ARFGEF1 antibody to recognize both protein products (Supplemental Fig. S2). Our results revealed no signal of the predicted protein size of ts-ARFGEF1 (142.3 kDa) and no difference in band patterns between before and after ts-ARFGEF1 knockdown (Supplemental Fig. S2), suggesting that ts-ARFGEF1 had low/no protein-coding potential.

**Figure 4.**
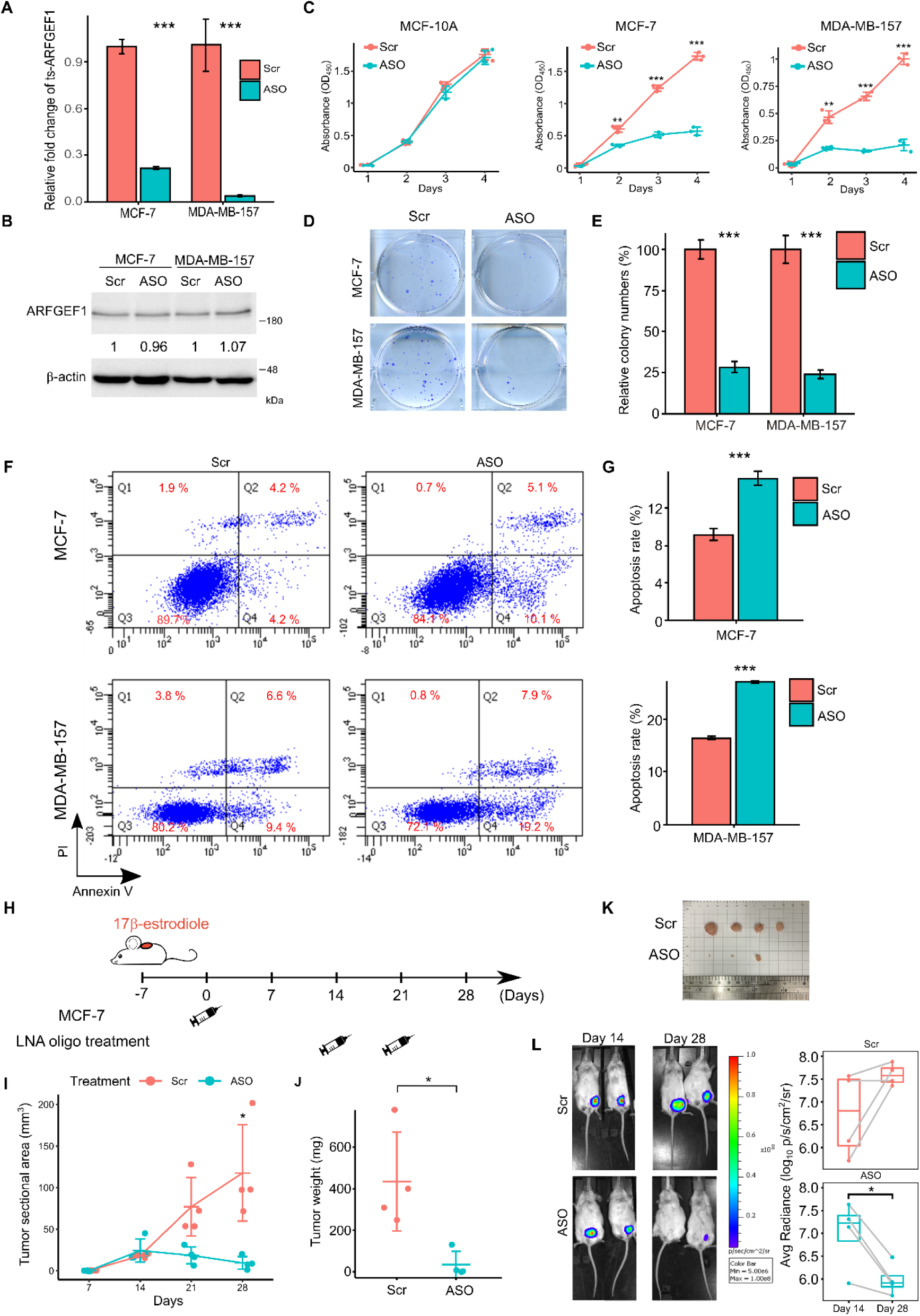
The biological role of ts-ARFGEF1 in breast cancer cell progression. **A** qRT-PCR analysis of the knockdown efficiency after transfecting MCF-7 and MDA-MB- 157 cells with the LNA ASO-mediated knockdown of ts-ARFGEF1. **B** Western blot analysis for the effect of ts-ARFGEF1 knockdown on the protein level of ARFGEF1. The relative expression of each protein was normalized to β-actin. **C-G** The effects of ts-ARFGEF1 knockdown on cell proliferation (via a CCK-8 kit; **C**), colony formation (via a clonogenicity assay; **D, E**), and apoptosis rate (via a flow cytometry assay; **F, G**) in MCF-7 and MDA-MB-157 cells. **H-L** The potential therapeutic effect of ts- ARFGEF1 on breast cancer growth *in vivo*. For **H**, a schematic diagram represented the workflow for examining the therapeutic effect of ts-ARFGEF1 knockdown in nude mice, in which nude mice were implanted with 17 β-estradiol in the shoulder area and then intraperitoneally implanted with scrambled control MCF-7 and ts-ARFGEF1 knockdown MCF-7 cells, respectively (n = 4 for each group). The tumor volume (**I**) and weight (**J**) of xenografts from MCF-7 cells with or without ts-ARFGEF1 silence in nude mice were examined. The representative morphological (**K**) and bioluminescent (**L**) images represented the nude mice from control or ts-ARFGEF1 knockdown groups. The tumor growth was evaluated by the quantification of average fluorescence radiance in each sample (**L, right**). Error bars in **A, C, E, G** were shown as the means ± standard deviation of three independent experiments. *P* values in **A, C, E, G, I, J** were determined using two-tailed *t*-test. *P* value in **L** was determined using paired two-tailed *t*-test. \**P* < 0.05, \*\**P* < 0.01, and \*\*\**P* < 0.001.

To determine the effects of ts-ARFGEF1 on tumor development *in vivo*, we orthotopically injected control MCF-7 and ts-ARFGEF1 knockdown MCF-7 cells into nude mice to establish a xenograft tumor model, respectively. Electronic caliper measurements revealed that the growth rates of the tumors derived from ts-ARFGEF1 knockdown were significantly lower than those derived from the control cells (Supplemental Fig. S3A). Consistently, we observed markedly lower tumor weights and smaller tumor volumes in the mice inoculated with ts- ARFGEF1-knockdown MCF-7 cells than in those inoculated with control cells (Supplemental Figs. S3B-D). To further examine whether ts-ARFGEF1 knockdown can serve as a therapeutic target for suppressing tumor growth, MCF-7 cells were first orthotopically injected into nude mice. Tumors were then allowed to grow for 14 days to reach a considerable size (∼25 mm). Next, these mice were treated with LNA ASOs through intraperitoneal injections (Fig. 4H). We observed that the treatment of LNA ASOs significantly reduced the growth rate of the tumors as compared with the control (Fig. 4I). At 28 days post-inoculation, the tumor volumes and weighs of the mice with the treatment of LNA ASOs were indeed greatly repressed (Fig. 4J-L). The above *in vitro* and *in vivo* experiments thus indicated that the disruption of ts-ARFGEF1 expression can inhibit tumor cell growth.

### Potential regulatory role of ts-ARFGEF1

Since ts-ARFGEF1 is more abundant in the nucleus than in the cytoplasm (Fig. 2C and Supplemental Fig. S1), we speculated that ts-ARFGEF1 may regulate the expression of the downstream genes at the transcriptional or post-transcriptional levels. We performed microarray analysis to examine the effect of ts-ARFGEF1 knockdown on the global gene expression profiling in MCF-7 cells. Gene set enrichment analysis (GSEA) based on the Hallmark gene sets from the molecular signature database (Subramanian et al. 2005) revealed that the top 10 upregulated and downregulated gene sets in ts-ARFGEF1 knockdown cells were associated with the cell stress/death signaling pathways (e.g., p53 signaling, apoptosis, hypoxia, UV response, and unfolded protein responses (UPR)) and the pro-proliferative signaling pathways (e.g., E2F targets and G2M checkpoint), respectively (Fig. 5A). On the basis of the microarray data, we also identified 167 upregulated genes and 298 downregulated genes in ts- ARFGEF1 knockdown MCF-7 cells (Fig. 5B, Supplemental Fig. S4A, and Supplemental Table S2). In concordance with the results of GSEA, the upregulated genes were significantly enriched in the gene sets related to the UPR and p53 pathways (Fig. 5C and Supplemental Table S2). p53 is a well-known tumor suppressor gene associated with apoptosis; and the p53- mediated pathway is known to be one of the major apoptosis signaling pathways (Hussain and Harris 1998). We thus examined the expression level of p53 protein after ts-ARFGEF1 knockdown. Indeed, we observed that the protein expression levels of both p53 and proteolytic cleavage of PARP, an important hallmark of apoptosis (Kaufmann et al. 1993), increased with increasing LNA ASO concentration (Fig. 5C). The expression of the *TP53* downstream genes related to the biological outcomes of cell cycle arrest, apoptosis, autophagy, and tumor microenvironment was also significantly increased after the knockdown (Fig. 5E and Supplemental Fig. S4B).

**Figure 5.**
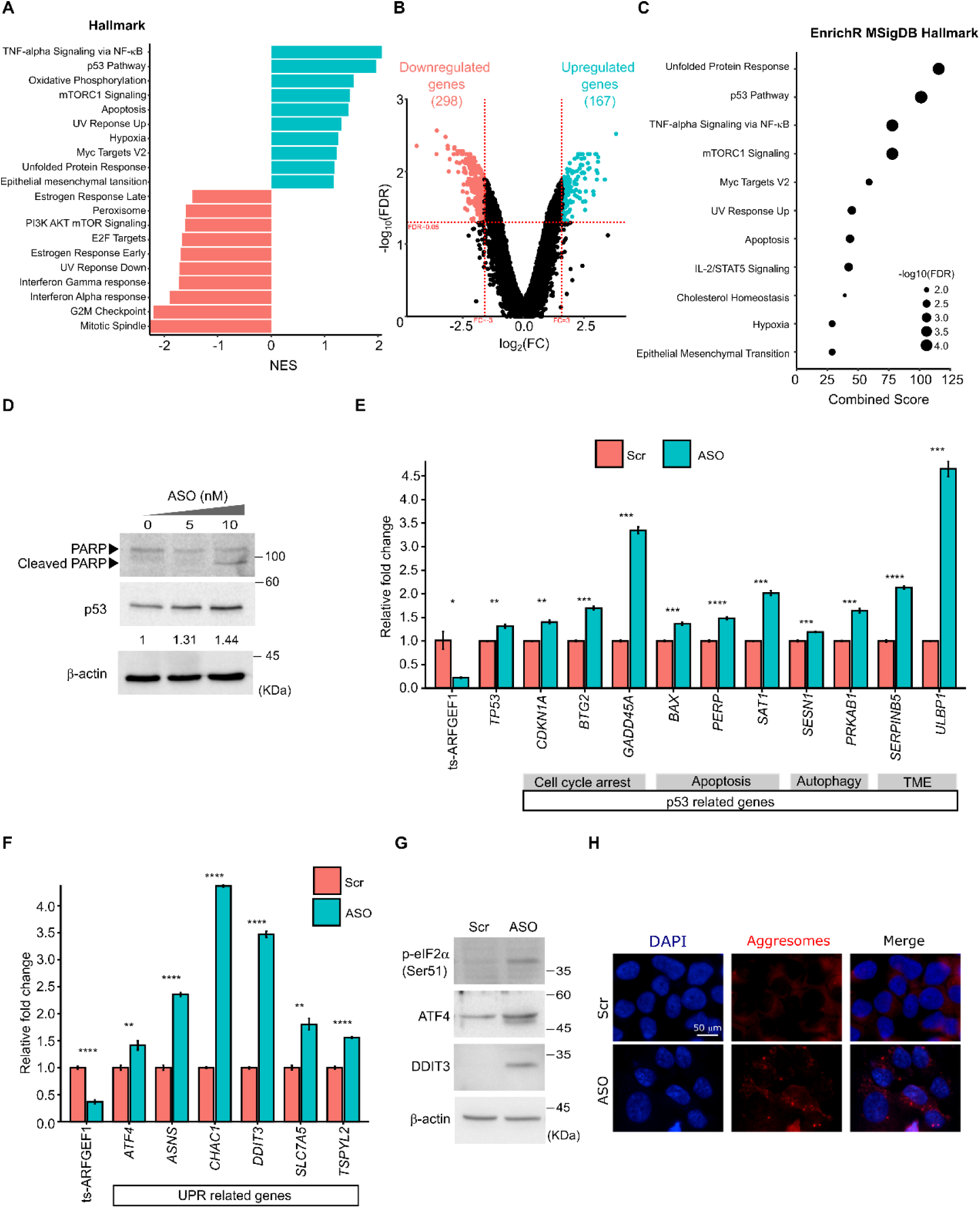
Transcriptome analysis of ts-ARFGEF1 knockdown reveals its potential role in ER homeostasis. **A** Gene set enrichment analysis (GSEA) of the microarray expression data from the ts-ARFGEF1 knockdown and scramble MCF-7 cells. The top 10 upregulated and downregulated Hallmark gene sets in ts-ARFGEF1 knockdown cells were shown. **B** Volcano plots representing the identified upregulated (167) and downregulated (298) genes in ts-ARFGEF1 knockdown cells with FDR<0.05 and |fold change|>3. **C** Gene enrichment analysis of the 167 upregulated genes on the basis of the Hallmark gene sets. **D** Western blot analysis of the correlation between the protein abundance of p53 and PARP cleavage products and the concentration of the LNA ASO- mediated ts-ARFGEF1 knockdown. **E** qRT-PCR analysis of the expression levels of *TP53* and the *TP53* downstream genes related to cell cycle arrest, apoptosis, autophagy, and tumor microenvironment before/after ts-ARFGEF1 knockdown in MCF-7 cells. **F** qRT-PCR analysis of the expression levels of the UPR-related genes before/after ts- ARFGEF1 knockdown. **G** Western blot analysis of the protein levels of phosphorylated eIF2α (p-eIF2α), ATF4, and CHOP before/after ts-ARFGEF1 knockdown. **H** The effect of ts-ARFGEF1 knockdown on the aggresome formation in MCF-7 cells. Immunostaining showed the aggresome formation (via an Abcam’s aggresome detection kit) in MCF-7 cells with or without ts-ARFGEF1 silence. Nuclei were stained with DAPI (blue). Error bars in **E, F** represented the means ± standard deviation of three independent experiments. *P* values were determined using two-tailed *t*-test. \**P* < 0.05, \*\**P* < 0.01, and \*\*\**P* < 0.001.

Meanwhile, the UPR was reported to be essential in regulating intracellular protein homeostasis(Hetz and Papa 2018). Under prolonged or strong endoplasmic reticulum (ER) stress, strong and sustained activation of the UPR can evoke apoptosis of cancer cells (Ishii et al. 2011; Hetz and Papa 2018). Therefore, we examined whether ts-ARFGEF1 played a role in regulating ER homeostasis and then leading to apoptosis. qRT-PCR analysis showed that the mRNA level of the key UPR genes (*TSPYL2* and *ATF4*) and the downstream effectors of *ATF4* (*DDIT3* (also known as CHOP), *ASNS*, *CHAC1*, and *SLC7A5*) were significantly elevated upon ts-ARFGEF1 knockdown (Fig. 5F and Supplemental Fig. S4B). It is known that activation of PERK/eIF2α/ATF4/CHOP pathway triggered by excessive ER stress can induce apoptosis(Hetz and Papa 2018). We thus examined whether ts-ARFGEF1 knockdown affected the PERK- mediated signaling pathway. Indeed, Western blot analysis showed that ts-ARFGEF1 knockdown caused a remarkable increase in the protein levels of phosphorylated eIF2α (p-eIF2α), ATF4, and CHOP (Fig. 5G). To further test whether ts-ARFGEF1 knockdown could elevate ER stress, we measured ER stress by monitoring the accumulation of misfolded/unfolded proteins. Indeed, ts-ARFGEF1 knockdown could contribute to aggresome formation in MCF-7 cells (Fig. 5H). Taken together, these results suggested that the knockdown of ts-ARFGEF1 could trigger unfolded protein response and thereby induce cell apoptosis by activating the PERK/eIF2a/ATF4/CHOP signaling pathway, implying the role of ts-ARFGEF1 in ER homeostasis in MCF-7 cells.

## Discussion

In this study, we focused on the identification and validation of intragenic ts-RNA events. We proposed a bioinformatics pipeline (“NCLscan-hybrid”) and multiple experimental validation steps to examine three types of potential false positives: (i) RT-based artifacts, (ii) circRNAs, and (iii) genetic rearrangements. For the bioinformatics steps, we employed the hybrid sequencing strategy with the integration of short and long RNA-seq reads and developed NCLscan-hybrid to detect intragenic ts-RNAs (Fig. 1A). There are several properties for this hybrid sequencing strategy while detecting intragenic NCL events. First, generally speaking, both read depth and sequencing quality are much lower in long-read sequencing data than in short RNA-seq reads. The ts-RNA candidates supported by both short and long reads should be more reliable than those supported by long reads only. Second, such a hybrid sequencing strategy can minimize the possibility of false positives derived from NGS platform specificity. Earlier studies had indicated a very low level of overlap between the NCL event identifications based on different NGS platforms (Maher et al. 2009; Wu et al. 2014). NGS platform-dependent ts-RNAs were suggested to be generated from experimental artifacts (Wu et al. 2014). With the hybrid sequencing strategy, NCLscan-hybrid may minimize the false positive from RT-based artifacts to a certain extent. Third, computational identification of intragenic NCL transcripts using short reads is often hampered by the difficulty of distinguishing between ts-RNA and circRNA isoforms. On the basis of TGS-based ultra-long RNA-seq reads, we proposed two features, out-of-circle and rolling circle, to distinguish between these two types of intragenic NCL transcripts. Here only the NCL transcripts supported by out-of-circle reads but not supported by rolling-circle reads were designated as ts-RNAs. We also utilized the WGS data of MCF-7 and did not find duplication or breakpoint events in the flanking intronic sequence of the identified ts-RNAs, minimizing the possibility of false positives from genetic rearrangements.

Correspondingly, we designed multiple experimental validation steps to examine each type of abovementioned false positives. For detection of RT-based artifacts, we respectively performed comparisons of different RTase products, RNA-FISH analysis, and RNase H cleavage to eliminate the possibility of RT-based artifacts for the identified ts-RNA candidate (Figs. 2B-2D). For discrimination between ts-RNA and circRNA forms, we first eliminated the possibility of the circRNA form by performing RNase R treatment (Fig. 2G). To confirm the existence of the ts-RNA form, we provided three lines of evidence. First, the ts-RNA candidate was a linear form based on qRT-PCR analysis with an oligo dT pull-down experiment (Fig. 2H). Second, we pulled down the selected ts-RNA candidate and sequenced the targeted transcript using the nanopore sequencer (Fig. 2I). We again demonstrated that the NCL candidate was supported by out-of-circle reads but not supported by rolling-circle reads (Fig. 2K). Third, RT-PCR experiments with multiple pairs of convergent PCR primers also showed that the NCL transcript spanned outside the potentially predicted circle sequence (Fig. 2L). For detection of genetic rearrangements, we performed PCR-Sanger sequencing and long PCR-nanopore sequencing to examine the existence of duplications/large insertions in the short and long introns flanked by the NCL junction, respectively (Figs. 2E and 2F).

We particularly emphasized the importance of discrimination between ts-RNA and circRNA forms for functional analysis of the selected NCL events. Generally, most detected intragenic NCL junctions are derived from *cis*-backsplicing; circRNAs are more abundantly expressed than ts-RNAs(Chuang et al. 2018). However, a considerable proportion of detected NCL junctions may be originated from *cis*-backsplicing and *trans*-splicing simultaneously (Yu et al. 2014; Chuang et al. 2018). These two types of NCL isoforms were observed to exhibit quite different expression patterns, suggesting their different regulatory roles (Chuang et al. 2018). If an observed NCL junction arises from both ts-RNA and circRNA products, it would be difficult to experimentally examine the biological functions of this NCL event through knockdown of the NCL junction because we could not determine whether the effect of the knockdown is derived from ts-RNA form or circRNA form. Therefore, it is necessary to definitely determine the type of NCL transcripts (ts-RNA or circRNA forms) for the examined NCL events before carrying out the subsequent functional analyses.

Regarding the identification of intragenic ts-RNAs, there are two inherent limitations for the strategy based on out-of-circle long reads, while (i) the predicted exonic regions of the examined ts-RNAs and the corresponding circRNA products completely overlap with each other and (ii) the same exonic region is repeated more than two times in a ts-RNA product. For the first scenario, a ts-RNA is formed by two separate precursor mRNAs from the same gene, in which all exons of the corresponding co-linear host gene are included in the ts-RNA isoform. For example, the host gene is a single exon gene (or see Case 1 illustrated in Addition file: Fig. S5). In such cases, NCLscan-hybrid is not destined to find any out-of-circle reads to support this ts-RNA form. For the second scenario, the same exonic region is repeated more than two times in a ts-RNA product (e.g., Case 2 in Addition file: Fig. S5). In this case, a ts-RNA is formed by more than two separate precursor mRNAs from the same gene. Such an NCL event may be misjudged as a circRNA because long reads (i.e., rolling-circle reads) may capture the whole exonic region of the ts-RNA and cross the NCL junction twice (or more than two times). Extremely, like the case illustrated in Case 3 of Addition file: Figure. S5, NCLscan-hybrid may simultaneously find out-of-circle reads and rolling-circle reads to support this NCL event. For these scenarios, further experimental validation steps (such as the abovementioned validations) can be applied to discrimination between intragenic ts-RNAs and circRNAs.

Upon multiple steps of bioinformatics-based screening processes and experimental validations, we identified and carefully confirmed an intragenic ts-RNA, ts-ARFGEF1, in breast cancer cells. Subsequent experiments demonstrated that the disruption of ts-ARFGEF1 expression can significantly affect breast cancer cell proliferation, colony formation, and apoptosis rate (Figs. 4C-4G). ts-ARFGEF1 silencing can significantly retard tumor cell growth in a breast cancer orthotopic mouse model (Supplemental Fig. S3 and Figs. 4H-4L). Our results further revealed the high association between ts-ARFGEF1 expression and key genes related to p53 and UPR signaling pathways (Figs. 5A-5C). The knockdown of ts-ARFGEF1 can activate the PERK/eIF2α/ATF4/CHOP pathway of UPR and the p53 pathway and accelerate aggresome formation in MCF-7 cells (Figs. 5D-5G), suggesting the regulatory role of ts- ARFGEF1 in ER homeostasis. It is known that UPR can act as a double-edged sword in responsible for cancer cell fate determined by the integration of pro-death and pro-survival signals (Sisinni et al. 2019). Activation of PERK/eIF2α/ATF4/CHOP pathway triggered by excessive ER stress can contribute to the induction of apoptosis through mediating p53 (Hetz and Papa 2018). Accumulating evidence has indicated the regulatory significance of non-coding RNAs (e.g., microRNAs, long non-coding RNAs, and circular RNAs) in ER stress response and cancer development(Zhao et al. 2020). Here we reported a new case of non-coding RNAs (ts-ARFGEF1) that can regulate ER homeostasis. It is worthwhile to further investigate the underlying molecular mechanisms of ts-ARFGEF1 in breast cancer. Of note, while we have demonstrated that ts-ARFGEF1 can regulate p53-mediated apoptosis and such a regulatory effect is not due to its cognate co-linear form (ARFGEF1), there is a technical obstacle to validate whether ARFGEF1 also acts as the similar regulatory role. It is challenging to disrupt *ARFGEF1* expression exclusively, because the exonic sequences of *ARFGEF1* overlaps with those of ts-ARFGEF1. Meanwhile, overexpression of *ARFGEF1* expression is also technically difficult due to the ultra-long coding sequence of *ARFGEF1* (5,547 bp). We thus suggest that functional analysis of co-linear RNAs should take into consideration the effect of ts-RNA forms if both co-linear RNA and ts-RNA forms of a gene are expressed in an examined sample.

In terms of the biogenesis of *trans*-splicing, we validated that RCS pairs in the introns flanking the NCL junction can promote ts-RNA formation. On the basis of ectopic expression experiment (Figs. 3A and 3B) and CRISPR-based endogenous genome modification experiments (Figs. 3C-3F), we successfully identified and confirmed an RCS pair that mediated ts-ARFGEF1 formation. Importantly, mutating nucleotide sequence at the RCS with CRISPR-based editing/deletion significantly repressed ts-ARFGEF1 expression without apparent effect on the protein expression of its cognate co-linear form (i.e., ARFGEF1) (Fig. 3G). Like the effect of ts-ARFGEF1 knockdown on cell proliferation in MCF-7 cells, the CRISPR-based endogenous genome modification indeed suppressed cell proliferative ability significantly (Supplemental Fig. S6A). We further found that the RCS-edited cells showed a significant increase in doxorubicin sensitivity (Supplemental Fig. S6B). Our results thus confirmed the requirement of the RCS pair for ts-ARFGEF1 formation and provided an efficient and specific approach to endogenous ts-RNA knockout at the genomic level.

In summary, this study systematically designed a bioinformatics pipeline based on a hybrid sequencing strategy and the corresponding experimental validation steps to minimize potential false positives from RT-based artifacts, circRNAs, and genetic rearrangements while detecting intragenic ts-RNAs. We thereby identified and confirmed an intragenic ts-RNA (ts-ARFGEF1) and demonstrated its regulatory role in p53-mediated apoptosis via PERK/eIF2α/ATF4/DDIT3 signaling pathway in breast cancer cells. In addition, for the first time, we provided the endogenous evidence to support ts-RNA biogenesis regulated by RCSs flanking NCL junctions. With the ability of discrimination between intragenic ts-RNAs and circRNAs, the established computational pipeline and validation steps are also applicable to the circRNA study. Our study thus describes a useful framework for rigorous characterization of transcriptionally non-co- linear RNAs, expanding the discovery of this important but understudied class of RNAs.

## Methods

### The NCLscan-hybrid pipeline

NCLscan-hybrid employed a hybrid sequencing strategy with integration of short and long RNA-seq reads to identify ts-RNAs (Fig. 1A). NCLscan (Chuang et al. 2016) was first utilized to search for all possible intragenic ts-RNA candidates according to the short reads generated from poly(A)-selected RNAs (Table 1). Genome assembly GRCh38 and Ensembl gene annotation (version 87) were used. For simplicity, this study focused on ts-RNAs derived from topologically distinct genomic loci only. The cases of ts-RNAs that involved co-linear exons arising from different alleles or opposite strands of the same gene loci (Gingeras 2009; Yu et al. 2014) were not considered. NCLscan-hybrid then generated a pseudo sequence by concatenating the exonic sequences flanking the NCL junction for each detected NCL event (within -100 nucleotides of donor site to +100 nucleotides of acceptor site) and aligned all TGS-based long reads against the 200 bp pseudo sequence using minimap2 (Li 2018) (Fig. 1B). A concatenated pseudo sequence may be shorter than 200 bp if one or both of the flanking exonic sequences are shorter than 100 bp. The minimap2 command lines were minimap2 -x map-pb - c --secondary=no for PacBio reads and minimap2 -x map-ont -c --secondary=no for nanopore reads, respectively. A long read was regarded to be a support of an NCL junction if the long read matched to ≥ 80% of the pseudo sequence with mapping quality = 60 and spanned the NCL junction boundary by ≥ 10 bp on both sides of the NCL junction. Only the long reads that supported an NCL junction were retained. After that, NCLscan-hybrid counted how many times the alignment of the long read crossed the NCL junction. Accordingly, each retained long read was then split into two or more segments at the NCL junction. Each segment was aligned against the reference genome. To minimize false positives from an alternative co-linear explanation or multiple hits, only the NCL events were retained if there were at least one long read(s) that satisfied the two rules simultaneously (Fig. 1B). First, all split segments of a supporting long read were uniquely matched (mapping quality=60) to the same gene loci using minimap2 -ax splice. Second, the integration of mapped segments can span the specified NCL junction. If a supporting long read had at least one segment(s) that mapped outside the potentially predicted circle sequences of the NCL event, the long read was designated as an “out-of-circle read”. An NCL event supported by at least one out-of-circle read(s) was regarded as a ts-RNA. On the other hand, if a supporting long read was split into three or more segments at the NCL junction, in which no segment mapped outside the potentially predicted circle sequences of the NCL junction, the long read was designated as a “rolling-circle read”. An NCL event supported by at least one rolling-circle read(s) was regarded as a circRNA.

### Cell Culture

MCF-7 cells were cultured in Dulbecco’s modified Eagle’s medium (DMEM) supplemented with 10 % FBS and 1 % penicillin-streptomycin (Gibco, Cat. 15140-122) at 37°C and placed in a humidified atmosphere containing 5% CO_2_. Normal breast MCF-10A cells were culture in DMEM medium supplemented with 10% FBS, epidermal growth factor (20 ng/ml), insulin (0.01 mg/ml), hydrocortisone (500 ng/ml) and cholera toxin (100 ng/ml) under standard culture condition (37°C, 5% CO_2_). For MDA-MB-157 culture, the cells grew in Leibovitz’s L-15 medium (Gibco, Cat.11415-064) supplemented with 10 % FBS and 1 % penicillin-streptomycin and were maintained at 37°C. MCF-7 and MCF7-10A cell lines were a kind gift from Dr. Michael Hsiao. MDA-MB-157 cell line was a gift from Dr. Chun-Mei Hu’s lab.

### RNA-Fluorescence *in situ* hybridization (RNA-FISH)

The RNA-FISH experiment was conducted according to the manufacturer’s instruction (Stellaris RNA FISH protocol for Adherent Cells). In brief, MCF-7 cells were cultured in a 12- well plate (4 × 10^4^ cells/well) for two days of culture. Next, the cell was fixed with fixation buffer and permeabilized with 70% ethanol overnight. After the washing step, the cells were covered by a hybridization buffer containing 25 nM Cy3-labeled probes antisense targeting the NCL junction of ts-ARFGEF1 and incubated in the dark at 42 °C for 16 hours. The slides were washed with Wash buffer A and mounted onto glass slides using DAPI-Fluoromount-G mounting medium (SouthernBiotech, Cat. 0100-20).

### RNase H treatment

Total RNA was heated at 65 °C for 5 min in the presence of probes against the NCL junction and then annealed at room temperatures. The reactions were treated with RNase H (NEB, Cat. M0297S) in the provided reaction buffer for 5 min at 37 °C. The remaining RNA was reverse transcribed and quantified by qRT-PCR.

### qRT-PCR

Total RNA was extracted from cells with TRIzol Reagent (Life Technologies, Cat. 15596018) and the PureLink RNA Mini Kit (Thermo Fisher Scientific, Cat. 12183018A). Total RNA was reverse transcribed by SuperScript IV Reverse Transcriptase kit (Thermo Fisher Scientific, Cat. 18090010) primed with random hexamers and oligo(dT) primers. qRT-PCR was conducted on a QuantStudio 5 Real-Time PCR System (Applied Biosystems) using Luminaris Color HiGreen qPCR Master Mix (Thermo Fisher Scientific, Cat. K0391). *GAPDH* was used as an endogenous control for quantification. The relative expression levels were calculated by the 2^-ΔΔ^CT method.

### RNase R treatment

For the RNase R experiment, 1 μg total RNA was treated with or without RNase R (3U/μg total RNA, Lucigen, Cat. RNR07250) for 30 min at 37 °C to deplete linear RNAs. The products were reverse transcribed by Superscript IV VILO (Thermo Fisher Scientific, Cat. 11756050). After reverse transcription, the cDNA was quantified by qRT-PCR.

### Oligo-dT pull down

For the selection of poly(A) RNA, the pull-down assay was performed by the Oligotex mRNA kit (QIAGEN, Cat. 780022). The selected RNA was reverse transcribed by SuperScript IV Reverse Transcriptase kit (Thermo Fisher Scientific, Cat. 18090010) primed with oligo(dT) primers. After reverse transcription, the cDNA was quantified by qRT-PCR.

### TERRA-capture RNA

To capture the transcript of ts-ARFGEF1, we performed a TERRA-capture RNA experiment as previously described (Chu et al. 2017). In brief, 20 μg of Trizol-purified total RNA was treated with 4 U of TURBO DNase (Thermo Fisher Scientific, Cat. AM2238) at 37 °C 10 min in 100 μl with 1 μl RNaseOUT (Thermo Fisher Scientific, Cat. 10777019). The RNA was purified by Trizol. Next, 20 μg RNA was mixed with 10 pmol of biotin-labeled oligo probes against the NCL junction of ts-ARFGEF1 (TCCAAGCCAACTGTTCCGTT/3’BioTEG) or scramble negative control probe (GAATTCGTCCTCCGATTCAC/3’BioTEG) and denatured at 70 °C in 100 μl 6×SSC hybridization buffer for 10 min. We then lowered the incubation temperature to 44 °C (0.5 °C/s). RNA was captured by 100 μl Dynabeads MyOne Streptavidin C1 (ThermoFisher, Cat. 65001), washed with 2×SSC/0.1% NP40 at 37 °C 5 min for 4 times, washed twice with 1×SSC/0.1% NP40 at 37 °C, and rinsed once with 1×SSC at room temperature. After a series of wash steps, the captured RNA was eluted by 30 μl DEPC treated water at 70 °C for 5 min. Finally, 3 μl elution was used for the analysis of capture efficiency.

### Nanopore sequencing

After capturing the transcript of ts-ARFGEF1 by biotin-labeled probe against the NCL junction, captured RNA was used for generating cDNA for nanopore sequencing according to the ONT cDNA-PCR sequencing protocol by the manufacturers (Oxford Nanopore Technologies, SQK- PCS109). Briefly, complementary strand synthesis from RNA and strand switching was conducted by using kit-supplied oligonucleotides. Double strand cDNA was then generated by PCR amplification using primers containing the 5’ tags, facilitating the ligase-free attachment of Rapid Sequencing Adapters. Finally, three libraries with ∼11 ng PCR products were sequenced by an R9.4 Flow cell with three times priming. To examine the full-length sequence of Intron 19 of *ARFGEF1*, we conducted the long-range PCR experiment by PrimeSTAR GXL DNA polymerase (TaKaRa, Cat. R050B). A total of 100 fmol of gel-purified PCR products was sequenced by the protocol of ONT Amplicons by Ligation (Oxford Nanopore Technologies, SQK-LSK109). Briefly, DNA was repaired by FFPE (New England Biolabs, Cat. M6630) and end-prepped/dA-tailed by using the NEBNext End Repair/dA-Tailing Module (New England Biolabs, Cat. E7546). Next, the kit-supplied sequencing adapters were ligated onto the prepared ends. After purification by AMPure XP beads (Beckman Coulter, Cat. A63881), a total of 50 fmole DNA was sequenced by an R9.4 Flow cell.

### Biotin-labeled oligonucleotides for targeting NCL junctions

We pulled down the transcripts containing the target region directly and performed nanopore long-read sequencing for the RT-PCR amplicons. NCLscan-hybrid was performed to examine the generated nanopore reads. A total of 133 nanopore reads were observed to span the NCL junction of ts-ARFGEF1, of which 88 (66%) and 0 reads were determined as out-of-circle and rolling-circle reads, respectively.

### Detection of genetic rearrangements

To examine whether the identified NCL junctions were derived from genetic rearrangements, we downloaded the WGS data of MCF-7 (Li et al. 2016) (Table 1) from the NCBI GEO database at https://www.ncbi.nlm.nih.gov/sra/ (accession number SRX1705314). We then performed INTEGRATE (Zhang et al. 2016) with default parameters to identify structural variants in the MCF-7 genome and check whether duplication or breakpoint events occurred in the flanking introns of the identified NCL junctions. For ts-ARFGEF1, we further used the ONT MinION sequencer to sequence the genomic sequence of Intron 19 of *ARFGEF1* from MCF-7 cells. A total of 3,040,649 nanopore long reads were generated and then aligned against the reference genome (GRCh38) using minimap2. We found that 92 % (2,808,975/3,040,649) of the nanopore reads uniquely mapped (mapping quality=60) to Intron 19. The uniquely mapped reads were assembled by minimap2 (minimap2 -x ava-ont) and then formed a consensus sequence by Racon (Vaser et al. 2017). We aligned the consensus sequence against the genomic sequence including Exon 19, Intron19, and Exon20 (chr8:67,240,164-67,251,450) using MUMmer (version 3.0) (Kurtz et al. 2004) and generated the dot plot of alignment (Fig. 2F) using MUMmerplot (Kurtz et al. 2004).

### Minigene plasmid construction

The genomic segments containing Exon26-Intron26-Exon27 of *ARFGEF1* or Exon26-Intron26 (mutant)-Exon27 were generated by PCR amplification. The PCR products were inserted into the pCMV-FLAG vector by NEBuilder HiFi DNA Assembly Cloning Kit (NEB, Cat. E5520S). For propagation of the construct, NEB Stable Competent E. coli (NEB, Cat. C3040H) were transformed by adding 2 μl assembly products to 50 μl of cells according to the manufacturer’s protocol. To identify clones with correct inserts in the vector, 0.5 μg of each plasmid DNA was treated with 3U of HindIII (NEB, Cat. R3104) and XmaI (NEB, Cat. R0180S) for 1 hour at 37 °C and visualized on a 1 % agarose gel. The plasmids with the correct size of the insert were validated by Sanger sequencing.

### CRISPR-based endogenous genome modification experiments

The guided sequences (agaaagggggaaaaactga) for *ARFGEF1* Intron 26 base editing were cloned into a pLAS-CRISPR-ABEmax-NG vector; and the guided sequences (gagctttcttaaacaaaggg) for *ARFGEF1* Intron 19 base deletion were cloned into pLAS-CRISPR- Hifi-NG. Both endogenous genome modification experiments were performed by National RNAi Core Facility, Academia Sinica, Taipei, Taiwan. MCF-7 cells were seeded into 6 well plates and transfected with CRISPR constructs by TransIT-X2 reagent (Mirus, Cat. MR- MIR6004). A limiting dilution was conducted for single-cell cloning three days after puromycin selection. The selected clones were screened by PCR and sequenced by Sanger Sequencing. The editing level was calculated by the EditR algorithm (Kluesner et al. 2018). CRISPR-mediated indels were detected by CRISP-ID (Dehairs et al. 2016).

### Cell proliferation

CCK-8 kit was used to evaluate the proliferation of MCF-10A, MCF-7, and MDA-MB-157 cells. Transfected cells (3×10^3^ cells in culture medium) were laid into 96-well plates. After incubation for 24, 48, and 72 hours, 10 μl CCK-8 reagent was added to each well followed by incubation at 37 °C for 2 hours. Finally, the absorbance of each well was measured at 450 nm using a SpectraMax Paradigm Multi-Mode Detection Platform system (Molecular Devices).

### Colony formation assay

After 48 hours of the knockdown of ts-ARFGEF1 in MCF-7 cells, cells were collected, inoculated into a 6-well plate (300 cells/well), and cultured in an incubator with 5% CO_2_ at 37°C for 14 days. After two weeks of incubation, cells were washed with PBS, fixed with 4% paraformaldehyde, and stained with 0.05% crystal violet. Five fields were picked randomly. We then detected and photographed the colony formation under a camera and microscopy. The number of clones in each field was calculated using the ImageJ software (Schneider et al. 2012).

### Flow cytometry assay

The cells were inoculated into 6 wells (3×10^5^ cells in culture medium/well). After 2 days of transfection, the cells were trypsinized and stained with Annexin V Conjugates for Apoptosis Detection kit (Thermo Fisher Scientific, Cat. A13201). The apoptotic cells in each group were detected with the FACSCanto II flow cytometer (BD Biosciences). The experiment was repeated three times in each group.

### Protein extraction and western blotting

The cells were scraped off, pelleted, and lysed for 15 min on ice in RIPA buffer (Abcam, Cat. ab156034) supplemented with a complete protease inhibitor cocktail (Roche, Cat. 05892791001). The concentration of protein was adjusted using a BCA protein assay kit (Thermo Fisher Scientific, Cat. 23225). The protein samples were diluted in 4× loading buffer, heated at 95 °C for 5 min, and separated on SDS-PAGE. Proteins were transferred to the PVDF membrane using standard procedures. The membranes were blocked in 5% skimmed milk in TBST for 1 hour at room temperature. Next, the membranes were incubated at 4°C overnight with primary antibodies (Supplemental Table S3). After three times wash, the membranes were incubated with HRP-conjugated secondary antibodies (Supplemental Table S3) at room temperature for 1 hour. The membranes were detected using the ECL kit (PerkinElmer, Cat. NEL103E001EA) with the manufacturer’s protocol and the ChemiDoc imaging systems (Bio-Rad).

### Tumor xenografts and treatment with LNA-ASOs

The Institutional Animal Care and Utilization Committees of Academia Sinica approved all the animal experiments (IACUC no. 20-12-1573). Lentiviral Green fluorescent protein-luciferase (GFP-Luc) constructs were purchased from the National RNAi Core Facility Platform (Academia Sinica, Taiwan). Lentiviruses were produced by cotransfecting the shRNA-expressing vector with the pMDG and pΔ8.91 constructs into 293T cells using calcium phosphate. Viral supernatants were then harvested and used to infect MCF cells in the presence of 8 μg/mL polybrene (Santa Cruz Biotechnology). After GFP-Luc transfection, the cells will be sorted by flow cytometry to purify the MCF-7-Luc cell. The MCF-7-Luc cells (1×10^6^ cells in 100 μl) transfected with either scrambled or LNA modified antisense oligos against the NCL junction of ts-ARFGEF1 were orthotopically injected into 5–6-week NOD/SCID (NS) female mice (5 mice/group, total 10). The xenograft tumor proliferation was detected every week. The tumor volumes were calculated using the formula of length × width^2^. The tumor growth was detected using the IVIS imaging (IVIS, Xenogen) equipment. After 5 weeks of growth, the mice were sacrificed and the tumor was cut off for analysis. For investigation of the treatment effect of locked nucleic acid-modified antisense oligomers (LNA-ASOs) targeting the NCL junction of ts-ARFGEF1 on breast cancer growth, a pellet containing 0.72 mg of 17β-estradiol (60 days release, Innovative Research of America, Cat. SE121) was implanted into the shoulder area of NS mice (6 weeks) to promote MCF-7 tumor growth in a xenograft mouse model. After a week, 1×10^6^ MCF-7-Luc cells were orthotopically injected into the NS mice. Subsequently, administration of LNA-ASOs or LNA-scramble (10 mg/kg) to NS mice by intraperitoneal injection (4 mice/group, total 8). Tumor growth was monitored every week and detected using the IVIS imaging system. Tumor size was measured by an electronic caliper (length × width^2^). After 4 weeks of growth, the mice were sacrificed and the tumor was cut off for analysis.

### Microarray analysis

Total RNA of scrambled control MCF-7 and ts-ARFGEF1 knockdown MCF-7 cells was extracted with TRIzol Reagent (Life Technologies, Cat. 15596018). The experiments of RNA integrity evaluation (Agilent RNA 6000 Nanochip kit) and microarray hybridization (Clariom D human array) and the data collection were performed by the Affymetrix GeneChip System Service center in Genomics Research Center, Academia Sinica. On the basis of the microarray raw data, the differentially expressed genes (Supplemental Table S2) were identified using Transcriptome Analysis Console (TAC 4.0) software with FDR<0.05 and |fold change|>3. The signaling pathway analyses were performed by GSEA (Subramanian et al. 2005) and modEnrichr (Kuleshov et al. 2016; Kuleshov et al. 2019) using the Hallmark gene sets from the molecular signature database (Subramanian et al. 2005).

## Data access

NCLscan-hybrid is publicly accessible on Github at https://github.com/TreesLab/NCLscan-hybrid. The generated nanopore RNA-seq data for the pull-down transcripts containing the target region, nanopore long genomic reads for the sequence of *ARFGEF1* Intron 19, and microarray raw data with/without the treatment of ARFGEF1 knockdown in MCF-7 cells were deposited in the NCBI Gene Expression Omnibus (GEO) under accession numbers GSE197336, GSE197337, and GSE196514, respectively.

## Acknowledgement

This work is supported by both Genomics Research Center (GRC), Academia Sinica, Taiwan and the National Health Research Institutes, Taiwan (NHRI-EX110-11011B1) to T.-J.C. We thank National RNAi Core Facility, Academia Sinica, Taiwan and Affymetrix GeneChip System Service center in Genomics Research Center, Academia Sinica, Taiwan for performing microarray experiments and endogenous genome modification (editing and deletion) experiments, respectively. We also thank Prof. Chun-Mei Hu for providing the cell lines examined and Dr. Chien-Hsiu-Li for discussion.

## Authors’ contributions

T.-J.C. supervised and conceived this study. Y.-C.C. designed and performed experiments and participated in paper writing. C.-Y.C. developed NCLscan-hybrid and performed computational analyses. T.-W.C. debugged NCLscan-hybrid. M.-H.C. and M.H. provided NOD/SCID (NS) mice and designed tumor xenograft experiments in nude mice. H-M.K. and I.J.T. performed Oxford nanopore sequencing. T.-J.C. drafted and finalized the manuscript with input from Y.-C.C. and C.-Y.C. All authors read and approved the final manuscript.

